# *TAS2R38* predisposition to bitter taste associated with differential changes in vegetable intake in response to a community-based dietary intervention

**DOI:** 10.1101/249763

**Authors:** Larissa Calancie, Thomas C. Keyserling, Lindsey Smith-Taillie, Kimberly Robasky, Cam Patterson, Alice S. Ammerman, Jonathan C. Schisler

## Abstract

**Background:** Although vegetable consumption is associated with decreased risk for a variety of chronic diseases, few Americans meet the CDC recommendations for vegetable intake. The *TAS2R38* gene encodes a taste receptor that confers bitter taste sensing from chemicals found in some vegetables. Common polymorphisms in *TAS2R38*, including rs713598, rs1726866, and rs10246939, lead to coding substitutions that alter receptor function and result in the loss of bitter taste perception.

**Objective:** Our study examines whether bitter taste perception *TAS2R38* diplotypes were associated with vegetable consumption in participants enrolled in either an enhanced or a minimal nutrition counseling intervention within a community-based dietary intervention.

**Methods:** DNA was isolated from the peripheral blood cells of study participants (N = 497) and analyzed for polymorphisms using genotyping arrays. The Block Fruit and Vegetable screener was used to determine frequency of vegetable consumption. Mixed effects models were used to test differences in frequency of vegetable consumption between intervention and genotype groups over time.

**Results:** There was no association between baseline vegetable consumption frequency and the bitter taste diplotype (*p* = 0.937), however after six months of the intervention, we observed an interaction between bitter taste diplotypes and time (*p* = 0.046). Participants in the enhanced intervention increased their vegetable consumption frequency (*p* = 0.020) and within this intervention group, the non-bitter and intermediate-bitter tasting participants had the largest increase in vegetable consumption. In contrast, in the minimal intervention group, the bitter tasting participants reported a decrease in vegetable consumption.

**Conclusions:** Non‐ and intermediate-bitter taste blind participants increased vegetable consumption in either intervention group more than those who perceive bitterness. Future applications of precision medicine could consider genetic variation in bitter taste perception genes when designing dietary interventions.

**Author summary:** Most Americans under consume vegetables, despite clear associations between vegetable consumption and health benefits. Vegetables, such as broccoli, kale, and Brussels sprouts, contain bitter-tasting compounds, leading to taste aversion. Common polymorphisms on the *TAS2R38* taste receptor gene (rs713598, rs1726866, and rs10246939) influence the perception of bitter taste. We tested whether genetic predisposition to bitter taste influenced vegetable intake in a dietary intervention and found that *TAS2R38* diplotypes were related to vegetable consumption. Combining precision medicine approaches that identify taste profiles and personalizing dietary advice could help engage intervention participants and improve the impact of dietary interventions.

## Introduction

Few Americans consume the recommended amount of dark green and orange vegetables, despite the association between vegetable consumption and reduced risk of chronic diseases [1]. Public health practitioners and researchers aim to increase vegetable consumption through dietary interventions, but the impact of interventions on fruit and vegetable intake yields mixed results. For example, some interventions resulted in increased vegetable consumption by participants [2–4], whereas others did not significantly affect vegetable consumption [5]. In instances where interventions increase vegetable intake, the effects are generally small and participants often do not reach recommended intake levels [6,7].

One possible explanation for the mixed results of dietary intervention studies is heterogeneity of participants regarding characteristics that strongly influence vegetable intake, such as taste preferences. Taste is an important determinant of fruit and vegetable intake in adults and children in the United States (US) [8,9]. While phytonutrients in vegetables, such as phenols, flavonoids, isoflavones, terpenes, and glucosinolates, seem to be protective against certain cancers, their bitter taste can be a deterrent to consumption [10]. Vegetable sweetness and bitterness were found to be independent predictors of more or less preference for sampled vegetables and vegetable intake, respectively, and the ability to detect a bitter tasting compound called propylthiouricil (PROP) was related to vegetable taste preferences [11].

Identified in 2003 [12], the *TAS2R38* gene encodes a G protein coupled receptor that functions as a taste receptor, mediated by ligands such as PROP and phenylthiocarbamide that bind to the receptor and initiate signaling that can confers various degrees of taste perception [13]. Vegetables in the brassica family, such as collard greens, kale, broccoli, cabbage, and Brussels sprouts, contain glucosinolates and isothiocyanates, which resemble PROP, and therefore much of the perceived “bitterness” of these vegetables is mediated through *TAS2R38* [14]. Bitter taste receptors in the TS2R family are also found in gut mucosal and pancreatic cells in humans and rodents. These receptors influence release of hormones involved in appetite regulation, such as peptide YY and glucagon-like peptide-1, and therefore may influence caloric intake and the development of obesity [15]. Thus, bitter taste perception may affect dietary behaviors by influencing both taste preferences and metabolic hormonal regulation.

Three variants in the *TAS2R38* gene – rs713598, rs1726866, and rs10246939 – are in high linkage disequilibrium in most populations and result in amino acid coding changes that lead to a range of bitter taste perception phenotypes [16,17]. The PAV haplotype is dominant; therefore, individuals with at least one copy of the PAV allele perceive molecules in vegetables that resemble PROP as tasting bitter, and consequently may develop an aversion to bitter vegetables. In contrast, individuals with two AVI haplotypes are non-bitter tasters. PAV and AVI haloptypes are the most common, though other haplotypes exist that confer intermediate bitter taste sensitivity (AAI, AAV, AVV, and PVI) [18]. This taste aversion may apply to vegetables in general [19]. Therefore, dietary interventions aiming to increase vegetable intake may have different outcomes depending on individuals’ perceptions of the taste.

While many studies have examined whether certain participant and intervention characteristics influence differential response to dietary interventions, such as age, sex, race, education, disease state, and intervention delivery methods [20,21], we are not aware of studies examining whether genes associated with bitter taste perception moderate participants’ responses to dietary interventions. The Heart Healthy Lenoir (HHL) Project offers a unique opportunity to test a concept that the genetic predisposition to bitter taste perception may associate with a differential response to a dietary intervention among a diverse, community-based study population [22,23]. In this paper we tested the following two hypotheses:

1. Participants with the *TAS2R38* non-bitter taste diplotype will consume more servings of vegetables per day at baseline than participants with intermediate or bitter taster diplotypes.
2. The *TAS2R38* diplotype will moderate the effect of the HHL intervention on vegetable consumption such that participants with a bitter taste diplotype will have a lower increase in reported vegetables intake than the non-bitter taste participants after 6 months of the intervention.

## Results

### Study Population

#### Demographics

Participant characteristics at baseline and after 6-months are shown in Table 1. There were several differences between participants in the minimal versus the enhanced intervention groups. More women, Caucasians, highly educated, and non-smokers participated in the enhanced intervention compared to the minimal intervention at baseline. Despite attrition, there where were no significant differences in participant characteristics within each intervention group at baseline and after 6-months.

**Table 1:**
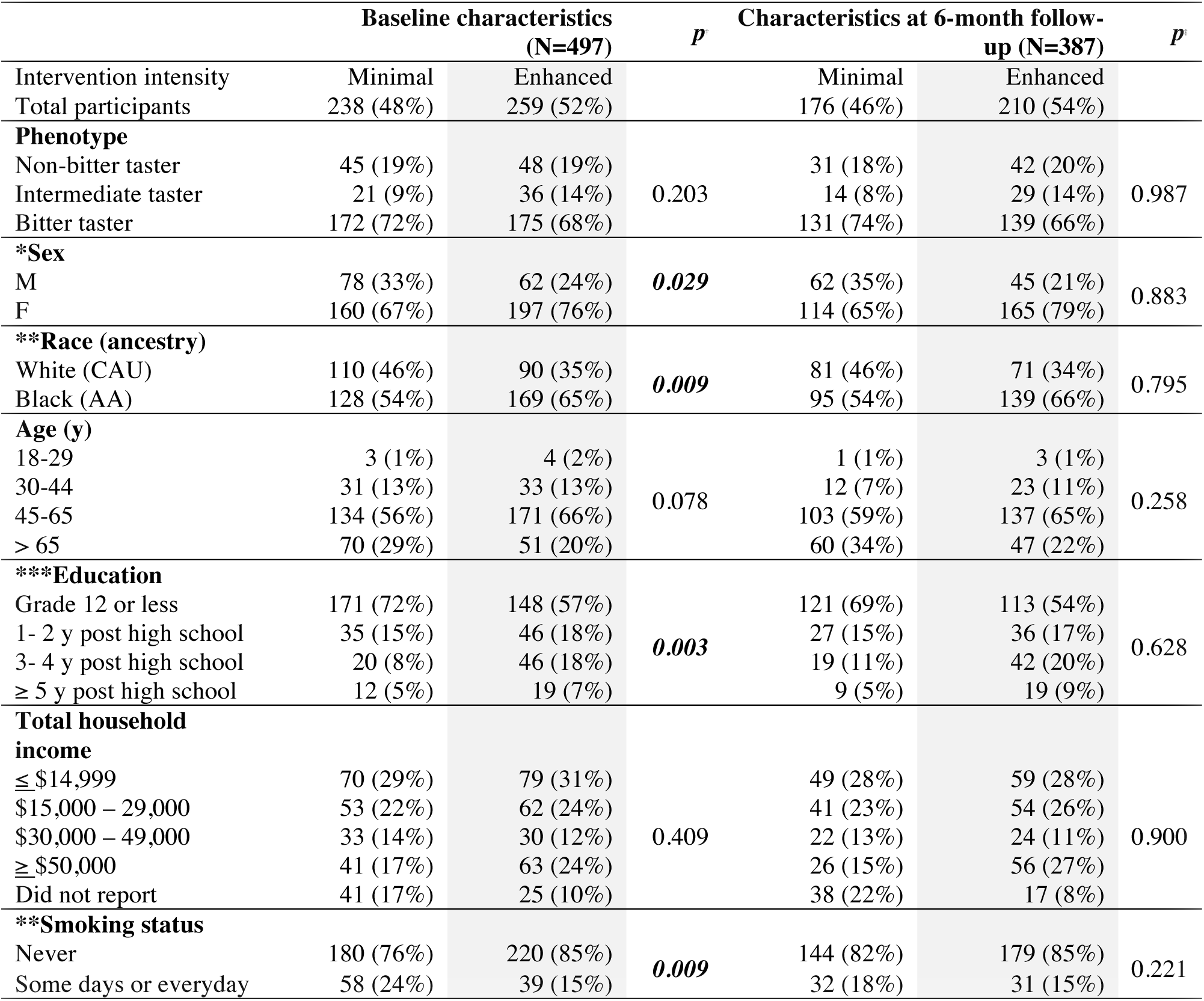
Study participant demographics at baseline and after 6-months of dietary intervention. Data presented as the frequency in each category for the indicated time point and intervention: *, **, and *** correspond to *p* < 0.05, < 0.01, or < 0.001 via a chi-squared comparing intervention intensity at baseline (†). The *p* value of a chi-squared test comparing baseline to 6-month follow-up is also indicated (‡).

| | Baseline characteristics<br>(N=497) | | $p$ | Characteristics at 6-month follow-up (N=387) | | $p$ |
| --- | --- | --- | --- | --- | --- | --- |
| Intervention intensity | Minimal | Enhanced |  | Minimal | Enhanced |  |
| Total participants | 238 (48%) | 259 (52%) |  | 176 (46%) | 210 (54%) |  |
| **Phenotype** |  |  |  |  |  |  |
| Non-bitter taster | 45 (19%) | 48 (19%) | 0.203 | 31 (18%) | 42 (20%) | 0.987 |
| Intermediate taster | 21 (9%) | 36 (14%) |  | 14 (8%) | 29 (14%) |  |
| Bitter taster | 172 (72%) | 175 (68%) |  | 131 (74%) | 139 (66%) |  |
| **\*Sex** |  |  |  |  |  |  |
| M | 78 (33%) | 62 (24%) | **0.029** | 62 (35%) | 45 (21%) | 0.883 |
| F | 160 (67%) | 197 (76%) |  | 114 (65%) | 165 (79%) |  |
| **\*\*Race (ancestry)** |  |  |  |  |  |  |
| White (CAU) | 110 (46%) | 90 (35%) | **0.009** | 81 (46%) | 71 (34%) | 0.795 |
| Black (AA) | 128 (54%) | 169 (65%) |  | 95 (54%) | 139 (66%) |  |
| **Age (y)** |  |  |  |  |  |  |
| 18-29 | 3 (1%) | 4 (2%) | 0.078 | 1 (1%) | 3 (1%) | 0.258 |
| 30-44 | 31 (13%) | 33 (13%) |  | 12 (7%) | 23 (11%) |  |
| 45-65 | 134 (56%) | 171 (66%) |  | 103 (59%) | 137 (65%) |  |
| > 65 | 70 (29%) | 51 (20%) |  | 60 (34%) | 47 (22%) |  |
| **\*\*\*Education** |  |  |  |  |  |  |
| Grade 12 or less | 171 (72%) | 148 (57%) | **0.003** | 121 (69%) | 113 (54%) | 0.628 |
| 1- 2 y post high school | 35 (15%) | 46 (18%) |  | 27 (15%) | 36 (17%) |  |
| 3- 4 y post high school | 20 (8%) | 46 (18%) |  | 19 (11%) | 42 (20%) |  |
| $\geq 5$ y post high school | 12 (5%) | 19 (7%) | | 9 (5%) | 19 (9%) | |
| **Total household income** |  |  |  |  |  |  |
| ≤ \$14,999 | 70 (29%) | 79 (31%) | 0.409 | 49 (28%) | 59 (28%) | 0.900 |
| \$15,000 – 29,000 | 53 (22%) | 62 (24%) | | 41 (23%) | 54 (26%) | |
| \$30,000 – 49,000 | 33 (14%) | 30 (12%) | | 22 (13%) | 24 (11%) | |
| ≥ \$50,000 | 41 (17%) | 63 (24%) | | 26 (15%) | 56 (27%) | |
| Did not report | 41 (17%) | 25 (10%) |  | 38 (22%) | 17 (8%) |  |
| **\*\*Smoking status** |  |  |  |  |  |  |
| Never | 180 (76%) | 220 (85%) | **0.009** | 144 (82%) | 179 (85%) | 0.221 |
| Some days or everyday | 58 (24%) | 39 (15%) |  | 32 (18%) | 31 (15%) |  |

#### TAS2R38 genetic characterization

All three alleles located in the *TAS2R38* gene are common variants in both African and Caucasian American populations [24] similar to our sample enrolled in HHL (**Supplemental Table 3**). In our CAU participants the three alleles had similar frequencies and were in high linkage disequilibrium (Table 2). The linkage disequilibrium was not as high across the pairwise allele comparisons in the AA participants (Rˆ2 range 0.46 – 0.95, D’ > 0.98) in part due to the difference in allele frequency of rs1726866 (Table 2). Therefore, we used the phased genotypes to determine the haplotypes found in our population. In our AA population, PAV was the most frequent haplotype, followed by AVI, haplotypes that encode the bitter and non-bitter polymorphisms, respectively (Table 2). This distribution was reversed in our CAU population. Demonstrating the genetic diversity between AA and CAU populations, nearly one-third the AA haplotypes were AAI (intermediate-taster phenotype) whereas the CAU haplotypes were almost exclusively PAV (bitter tasters) or AVI (non-bitter tasters) (96%).

**Table 2:** *TAS2R38* linkage disequilibrium and haplotype frequencies. Statistical analyses of linkage disequilibrium (LD) are represented by R-squared (Rˆ2), D, and Dprime values of the pairwise comparisons of the indicated SNPs from the AA and CAU participants. The plus strand haplotype sequence (HAPLO), the count of each haplotype, and the resulting amino acid sequence of the allele are indicated from the AA and CAU participants.

| **AA (N = 304)** |  |  |  |  |  |
| --- | --- | --- | --- | --- | --- |
| **LD analysis** | **SNP1** | **SNP2** | **$R^2$** | **D** | **Dprime** |
|  | rs10246939 | rs1726866 | 0.49 | -0.16 | -1.00 |
|  | rs10246939 | rs713598 | 0.95 | 0.24 | 0.99 |
|  | rs1726866 | rs713598 | 0.46 | -0.16 | -0.98 |
| **HAPLO** | C:G:G:307<br>PAV | T:A:C:190<br>AVI | T:G:C:104<br>AAI | C:G:C:5<br>AAV | T:A:G:2<br>PVI |
| **CAU (N = 201)** |  |  |  |  |  |
| **LD analysis** | **SNP1** | **SNP2** | **$R^2$** | **D** | **Dprime** |
|  | rs10246939 | rs1726866 | 0.98 | -0.25 | -0.99 |
|  | rs10246939 | rs713598 | 0.84 | 0.23 | 1.00 |
|  | rs1726866 | rs713598 | 0.84 | -0.23 | -1.00 |
| **HAPLO** | C:G:G:170<br>PAV | T:A:C:214<br>AVI | T:G:C:1<br>AAI | C:G:C:16<br>AAV | C:A:C:1<br>AVV |

The PAV is a dominant allele, therefore instead of relying on an index SNP or haplotypes, we used a dominant model to derive a bitter taste phenotype score based on the diplotype (Table 3). Contingency analysis of the bitter taste phenotype revealed that the percentage of bitter-tasting participants was similar between AA and CAU (Figure 1). However, among those not falling into the bitter tasting category, we observed a higher proportion of non-bitter tasters in CAUs (29%) versus AAs (12%) and three times as many intermediate tasters in AAs versus CAUs (Figure 1), likely due to the prevalence of the AAI (intermediate-taster) haplotype in our AA population (Table 2).

**Figure 1:**
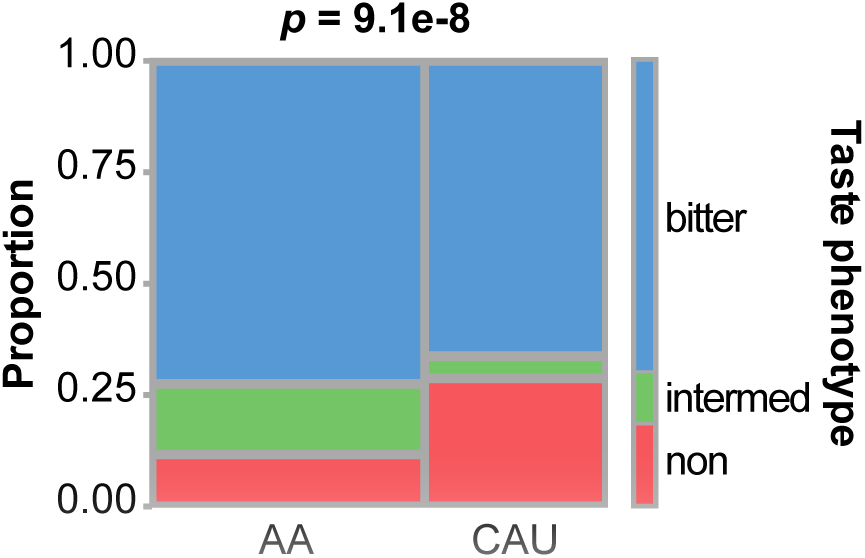
*TAS2R38* bitter taste phenotype distribution in the HHL cohort. Contingency plot and *p* value of the Fisher’s Exact Test in comparing the distribution (proportion) of taste phenotypes in the AA and CAU group.

**Table 3:** *TAS2R38* diplotype frequencies and associated phenotype. The distribution of diplotypes within the AA and CAU participants. The indicated bitter tasting phenotype for each diplotype is indicated.

| AA (N = 304) |  |  |
| --- | --- | --- |
| Diplotype | Freq | Phenotype |
| PAV / PAV | 0.286 | bitter |
| PAV / AVI | 0.270 | bitter |
| AAI / PAV | 0.155 | bitter |
| AVI / AVI | 0.118 | non |
| AAI / AVI | 0.115 | intermediate |
| AAI / AAI | 0.033 | intermediate |
| AAV / PAV | 0.013 | bitter |
| AVI / PVI | 0.007 | intermediate |
| AAV / AAI | 0.003 | intermediate |
| CAU (N = 201) |  |  |
| Diplotype | Freq | Phenotype |
| AVI / PAV | 0.438 | bitter |
| AVI / AVI | 0.289 | non |
| PAV / PAV | 0.184 | bitter |
| AAV / AVI | 0.040 | intermediate |
| AAV / PAV | 0.040 | bitter |
| AVI / AAI | 0.005 | intermediate |
| AVV / AVI | 0.005 | intermediate |

### Associations between vegetable consumption and genetic predisposition to bitter taste

#### Bitter taste diplotypes did not associate with differences in baseline vegetable intake

We first measured associations between baseline vegetable intake and *TAS2R38* phenotypes using model 1.Sex, education, and household income were positively associated with reported vegetable consumption frequency scores, as expected (Table 4). Participants reported similar vegetable consumption frequency independent of their genetic predisposition toward bitter taste sensitivity, *p* = 0.937 (Figure 2, Table 4). Thus, we rejected our first hypothesis that participants would report different vegetable consumption frequency scores at baseline according to their *TAS2R38* diplotype. These data suggest that within our HHL population, the *TAS2R38* polymorphisms were not associated with vegetable intake. This finding is consistent with another study examining the association between self-reported vegetable intake and PROP sensitivity in a community-based population [25].

**Figure 2:**
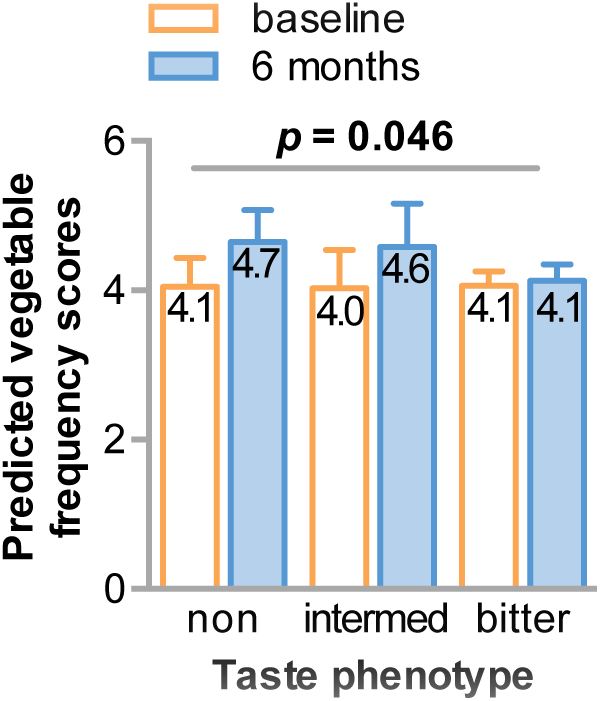
Vegetable intake at baseline and after 6-months categorized by *TAS2R38* bitter taste phenotype. Bar plots of the predicted vegetable intake adjusted for sex, ancestry, age, education, income, and smoking status represented by the mean ± 95% confidence intervals at either the onset of the study (baseline) or at the 6-month follow up, grouped by non-bitter (non), intermediate-bitter (inter), or bitter tasting phenotype. The *p* value of the interaction between taste phenotype and time is indicated.

**Table 4:** Regression coefficients for vegetable intake frequency at baseline (Model 1) and mixed effects coefficients at 6 months (Model 2). The coefficient of variation, standard error (SE), *t* statistic (Model 1), *z* score value (Model 2), 2-tailed *p* values (P > |*t*| or P > |*z*|), and 95% confidence intervals (CI) are provided: *, **, and *** correspond to *p* < 0.05, < 0.01, or < 0.001

**MODEL 1**
| Variables | Coefficient | SE | *t* | $P > \vert t\vert $ | 95% CI |
| --- | --- | --- | --- | --- | --- |
| Intermediate taster | -0.10 | 0.337 | -0.28 | 0.777 | -0.76 – 0.57 |
| Bitter taster | 0.01 | 0.231 | 0.03 | 0.979 | -0.45 – 0.46 |
| Non-smoker | 0.14 | 0.231 | 0.58 | 0.562 | -0.32 – 0.59 |
| **Female | 0.63 | 0.199 | 3.15 | **0.002** | 0.24 – 1.02 |
| Age | 0.01 | 0.008 | 1.88 | 0.061 | -0.001 – 0.03 |
| *Education | 0.08 | 0.037 | 2.08 | **0.038** | 0.004 – 0.15 |
| ***Income | 0.14 | 0.034 | 4.11 | **<0.001** | 0.07 – 0.21 |
| Race | 0.14 | 0.195 | 0.74 | 0.459 | -0.24 – 0.53 |
| Constant | 0.80 | 0.766 | 0.97 | 0.335 | -0.77 – 2.25 |

**MODEL 2**
| Variables | Coefficient | SE | *z* | $P > \vert z\vert $ | 95% CI |
| --- | --- | --- | --- | --- | --- |
| Intermediate taster | 0.06 | 0.501 | 0.13 | 0.899 | -0.92 – 1.05 |
| Bitter taster | 0.37 | 0.313 | 1.17 | 0.242 | -0.25 – 0.98 |
| Enhanced intervention group | 0.19 | 0.387 | 0.49 | 0.621 | -0.57 – 0.95 |
| Inter.: Enhanced | -0.17 | 0.644 | -0.27 | 0.791 | -1.43 – 1.09 |
| Taster: Enhanced | -0.70 | 0.434 | -1.54 | 0.123 | -1.52 – 0.18 |
| 6-month follow-up | 0.46 | 0.338 | 1.36 | 0.174 | -0.20 – 1.12 |
| Inter.: 6-months follow-up | -0.26 | 0.601 | -0.43 | 0.671 | -1.43 – 0.92 |
| Taster: 6-months follow-up | -0.89 | 0.376 | -2.38 | 0.018 | -1.63 – -0.16 |
| Enhanced: 6-months | 0.25 | 0.450 | 0.56 | 0.573 | -0.63 – 1.13 |
| Inter.: Enhanced: 6-month follow-up | 0.40 | 0.758 | 0.53 | 0.598 | -1.09 – 1.89 |
| Taster: Enhanced: 6-month follow-up | 0.68 | 0.505 | 1.35 | 0.177 | -0.31 – 1.67 |
| Non-smoker | 0.30 | 0.198 | 1.54 | 0.123 | -0.08 – 0.69 |
| ***Female | 0.70 | 0.166 | 4.22 | **<0.001** | 0.38 – 1.02 |
| Age | 0.01 | 0.007 | 1.55 | 0.122 | -0.003 – 0.02 |
| **Education | 0.09 | 0.031 | 2.89 | **0.004** | 0.03 – 0.15 |
| ***Income | 0.14 | 0.028 | 4.93 | **<0.001** | 0.08 – 0.19 |
| Race | -0.01 | 0.164 | -0.01 | 0.994 | -0.32 – 0.32 |
| Constant | 0.75 | 0.164 | 1.12 | 0.264 | -0.56 – 2.05 |

#### Participants with non-bitter or intermediate-bitter taste diplotypes increased vegetable intake after the intervention

Using model 2, we incorporated variables to measure the impact of the different interventions over time and to measure interactions between *TAS2R38* diplotypes, intervention intensity, and time (Table 4). We observed the same associations between reported vegetable consumption frequency scores and sex, education, and household income. Consistent with our second hypothesis, we observed an interaction between phenotype and time (Figure 2). Non-bitter taste participants reported 0.65 higher vegetable intake frequency scores, or about 0.20 servings of green salads or other vegetables per day, at the end of the intervention. Vegetable intake frequency scores also increased by 0.55 among intermediate bitter tasters. Intake scores only increased 0.04 among bitter tasters at the end of the intervention. Importantly, we did not see differences in participant demographics (Table 1) or allele frequencies, linkage disequilibrium, or haplotype distributions (**Supplemental Tables 3, 4, 5**) due to intervention attrition at the 6-month time point.

#### Vegetable intake increased in the enhanced dietary intervention

Given the enhanced intervention included tailored dietary goals and behavior change strategies, we hypothesized that participants in the enhanced intervention would have a greater increase in vegetable intake. As expected, the change in vegetable intake frequency scores was higher in the enhanced intervention group compared to the minimal group over time (Figure 3). In fact, participants in the minimal intervention group reported a decrease of 0.19 in vegetable intake frequency scores, whereas participants in the enhanced intervention group increased their reported scores by 0.58, suggesting that the enhanced intervention contributed to dietary changes regarding vegetable intake.

**Figure 3:**
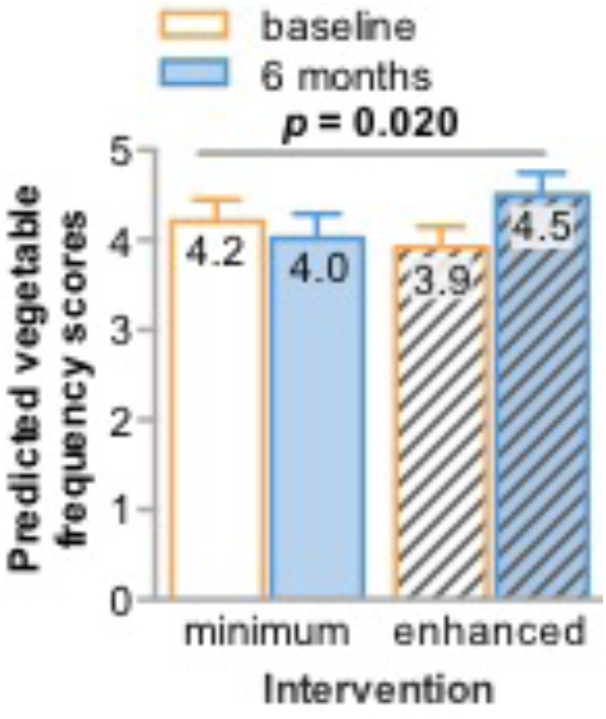
Vegetable intake at baseline and after 6-months categorized by intervention intensity. Bar plots of the predicted vegetable intake adjusted for sex, ancestry, age, education, income, and smoking status represented by the mean ± 95% confidence intervals at either the onset of the study (baseline) or at the 6-month follow up, grouped into the minimal or enhanced intervention group. The *p* value of the interaction between intervention intensity and time is indicated.

#### Bitter taste perception and the intensity of the dietary intervention may influence vegetable intake

Although the enhanced intervention associated with increased reported vegetable intake (Figure 3), could this response be modified by the *TAS2R38* phenotype? Despite significant main effects, the three-way interaction between intervention group, phenotype, and time was not statistically significant, *p* = 0.392. Still, the 3-way interaction analysis trends similar to those seen in the 2-way interactions (Figure 4). Non-bitter and intermediate-bitter tasting participants in the enhanced intervention increased their vegetable intake frequency score the most (delta = 0.71 and 0.89, respectively). Consistent with our hypothesis, bitter tasting participants in the minimal intervention were the only group that decreased their vegetable intake (delta = −0.44), however there was an increase among bitter tasting participants in the enhanced intervention (delta = 0.50). Our data suggest that these *TAS2R38* alleles and resulting phenotypes may impact a person’s response to dietary interventions regarding vegetable intake.

**Figure 4:**
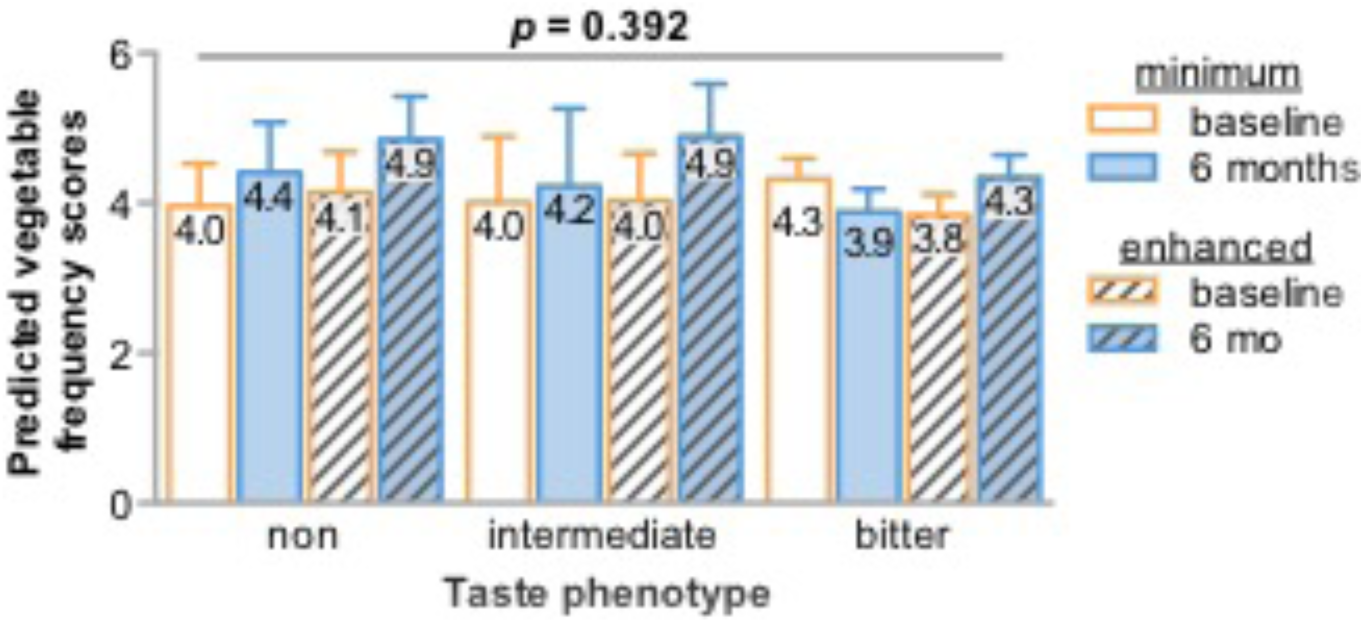
Vegetable intake at baseline and after 6-months in either intervention group categorized by *TAS2R38* bitter taste phenotype. Bar plots of the predicted vegetable intake adjusted for sex, ancestry, age, education, income, and smoking status represented by the mean ± 95% confidence intervals at either the onset of the study (baseline) or at the 6-month follow up, grouped by non-bitter (non), intermediate-bitter (inter), or bitter tasting phenotype within each intervention (minimum or enhanced). The *p* value of the three-way interaction between taste phenotype, time, and intervention intensity is indicated.

#### Vegetable intake associated specifically with *TAS2R38* variants and not other variants in related TAS2R genes

Other genes in *TAS2R* family are also implicated in taste perception, neuroendocrine function, appetite, and satiety [26] as well as human aging [27]. We extracted the genotypes of these related family members (**Supplemental Table 6**) and along with the *TAS2R38* variants we used principal components analysis with the adjusted predicted vegetable intake as a supplementary variable to determine if other *TAS2R* genes associate with the responsiveness to our dietary interventions. In our AA and CAU groups we identified the two components that corresponded to the highest loading for vegetable intake (Figure 5A, 5B). Not surprisingly, this resulted in segregation of the *TAS2R38* bitter taste phenotypes and revealed that the three *TAS2R38* alleles were highly correlated to the variance of PC4 or PC2 in the AA or CAU groups, respectively (**Supplemental Table 7**). We also identified another associated locus common to both AA and CAU populations that harbors *TAS2R20* and *TAS2R50* (Table 5, **Supplemental Table 7**). However, when we used a mixed model approach to look at the association of these individual SNP or the SNP: time interaction and reported vegetable intake, we only observed an association with two *TAS2R38* alleles, rs713598 and rs10246939. Another locus of interest included the *TAS2R3*, *TAS2R4*, and *TAS2R5* genes that had high correlation in PC2 in the CAU group (Figure 5B, **Supplemental Table 7**). However, like the other loci we analyzed, we did not find any association with vegetable intake either analyzed with both populations or only within the CAU group (**Supplemental Table 8**). These data suggest that *TAS2R38* is likely the largest genetic contributor to our association analysis. The other SNPs we identified in this analysis, however, may play other roles that contribute to taste perception and diet.

**Figure 5.**
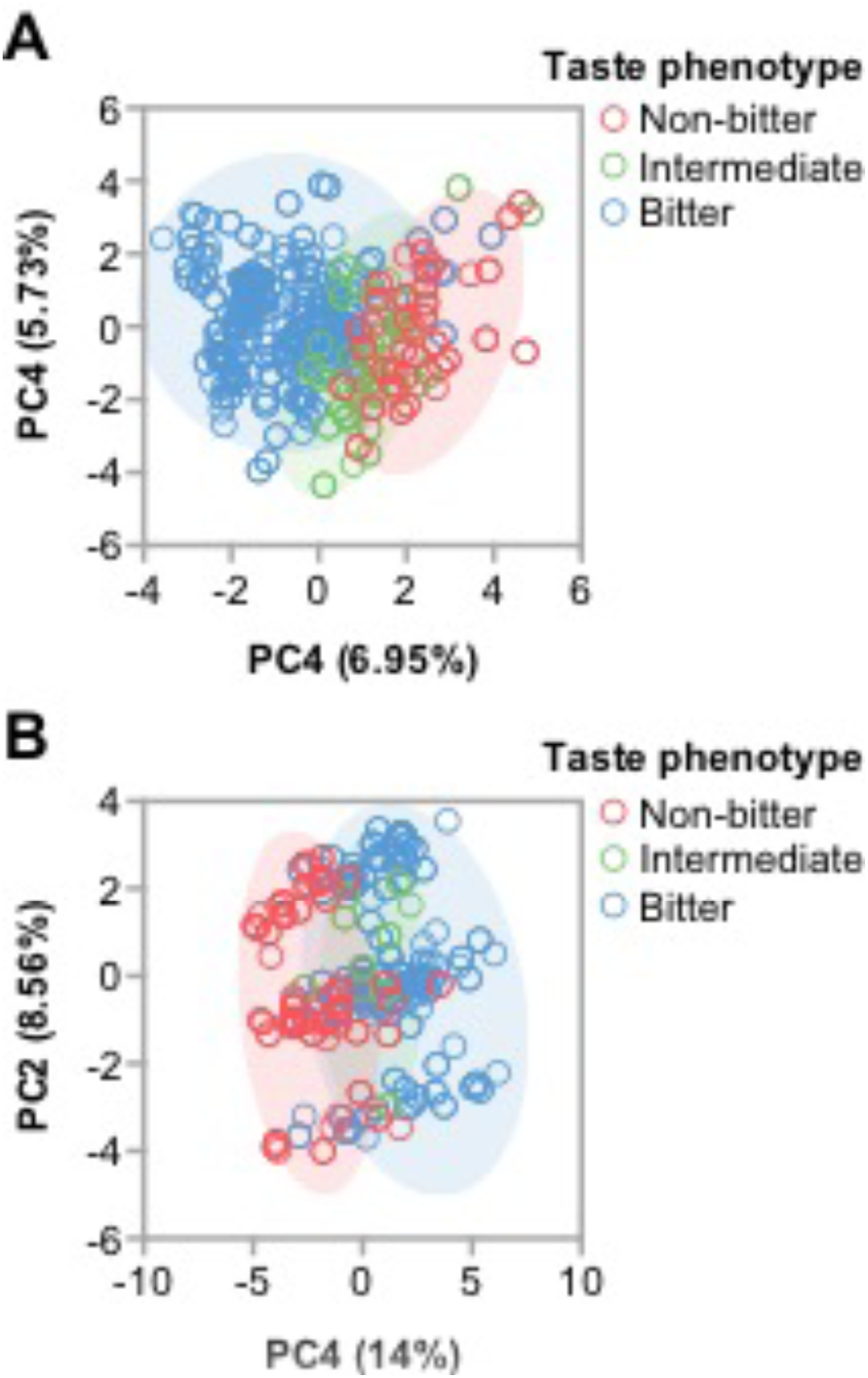
Multivariate analysis of *TAS2R* polymorphisms in the HHL cohort. Principal component scatter plots of (**A**) AA and (**B**) CAU groups colored by the *TAS2R38* phenotype. The percent variance explained by the indicated principal component (PC) is indicated.

**Table 5:** SNP and SNP-time associations with vegetable intake. The *p* values of the association of either the indicated SNP or the SNP: time interaction with reported vegetable intake. The location of the gene is indicated by chromosome (Chr) and position (Pos).

| SNP | Gene | Chr | Pos | p (SNP) | p (SNP:time) |
| --- | --- | --- | --- | --- | --- |
| **rs713598** | *TAS2R38* | 7 | 141673345 | 0.0659 | **0.0147** |
| **rs10246939** | *TAS2R38* | 7 | 141672604 | 0.0659 | **0.0147** |
| **rs1726866** | *TAS2R38* | 7 | 141672705 | 0.1208 | 0.1452 |
| **rs10772408** | *TAS2R49* | 12 | 11151599 | 0.4936 | 0.7443 |
| **rs1376251** | *TAS2R49* | 12 | 11138852 | 0.3534 | 0.9068 |
| **rs7301234** | *TAS2R49* | 12 | 11150884 | 0.2838 | 0.7276 |

## Discussion

The primary goal of HHL was to reduce CVD-related health disparities in a rural population in North Carolina. In this report, we tested the concept that participants in a dietary intervention designed to promote heart healthy eating patterns may respond differently according to their genetic predisposition of bitter taste perception mediated by the *TAS2R38* gene and allelic variants that can affect receptor signaling and hence, perception of bitter taste compounds found in many vegetables. Our HHL sample was represented by two ancestral populations, African and Caucasian Americans, and we were cognizant of the genetic population structure of our cohort. When we analyzed the diplotypes and corresponding phenotypes of our cohort, we observed similar proportion of bitter taste participants in the AA and CAU groups (Figure 1). There was a striking difference, however, in the proportion of non-bitter and intermediate bitter tasters such that the CAU group had nearly triple the frequency of non-bitter tasters (Figure 1), consistent with a recent study on the natural selection of *TAS2R38* haplotypes [24]. Although we lacked the power to stratify our HHL cohort for robust, focused analyses within each ancestry group, we accounted for ancestry in our analyses and the variable accounting for ancestry in either of our models did not approach our defined level of statistical significance (Table 4). Although these data suggest that ancestry did not associate with changes in reported vegetable consumption in our cohort, future studies should consider and seek to define differences in allele frequency and interactions with other biological factors that contribute to taste perception in distinct ancestral populations to determine the applicability of precision medicine to dietary interventions.

We found differences in vegetable consumption frequencies between intervention participants at follow-up according to their bitter taste perception phenotype characterized by common coding variants in the *TAS2R38* gene (Figure 2). Participants with *TAS2R38* diplotypes associated with non-bitter tasting increased vegetable consumption more than participants whose genotypes were associated with bitter taste perception (Figure 2). Our findings are consistent with other studies that observed differential vegetable preferences according to the presence of bitter taste perception SNPs [11,28]. However, other studies suggest that bitter taste sensitivity is not associated with food selection due to other factors such as attitudes toward foods, cultural norms, and one’s food environment [29,30]. More research is needed to better understand how genetic taste variation and other factors influence vegetable selection and consumption [30], and importantly, how this information can help inform dietary interventions.

Not surprisingly, we also found that participants in the enhanced dietary intervention increased their vegetable intake frequency scores more than those in the minimal intervention (Figure 3). A review of behavioral interventions aiming to increase vegetable intake found that 17 of 22 studies reported small, but significant increases in vegetable intake [21]. Many dietary intervention studies aim to change servings of total fruits and vegetables, while ours only examined a subset of vegetable intake (green salads and other vegetables) and likely explains the small changes we observed in daily servings of vegetables after the intervention. Moreover, the study participants reported very low intake of vegetables as baseline; in retrospect, participants may have benefitted from a more intensive vegetable consumption focus in the intervention than they received. In some cases, participants in the minimal intervention group reported lower vegetable intake frequency scores after 6 months than at baseline (Figure 3).

Participants who took part in the enhanced intervention increased their vegetable intake over the course of the intervention, irrespective of the *TAS2R38* phenotype, whereas participants in the minimal intervention showed mixed results based on *TAS2R38* phenotype (Figure 4). Non-bitter taste participants in the minimal intervention group increased their vegetable intake while bitter tasters in the same intervention group decreased their vegetable consumption (Figure 4). Our findings demonstrate that all participants in the enhanced condition, even those who are likely to perceive bitterness in some vegetables, increased vegetable consumption during the intervention. Biological sensitivity to bitter taste is likely one of many factors contributing to participants’ decisions about vegetable consumption. Participants that perceive bitterness may choose to consume vegetables that are less bitter, such as carrots or cooked vegetables [31] or food preparation strategies that minimize the bitter taste. Participants may have also modified their preferences toward vegetable consumption over the course of the enhanced intervention; studies suggest that repeated exposure to foods and beverages can alter preferences for those foods and beverages [32–34]. Since participants were receiving information about the benefits of a vegetable-rich diet, they may have been more willing to overcome taste aversions and perhaps even modify their taste preferences during the 6-month enhanced intervention.

There were several limitations in this study. Frequency of vegetable intake questions did not specifically target vegetables that are high in bitter compounds [11,31]. Additionally, cooking methods were not assessed, and cooking can affect consumers’ vegetable preferences [35,36]. Moreover, we did not include self-reported vegetable juice and vegetable soup intake in our outcome variable. These items were excluded because they are likely to have added salt or sugar, which suppresses bitterness [36,37]. Also, there was 22% attrition at the 6-month follow up; however, the haplotype frequencies were similar at baseline and follow-up (**Supplemental Table 5**, so the differences seen between baseline and 6 months are not likely due to differences in genotypes. Additionally, our sample size limited our ability to detect a statistically significant interaction between genotype and intervention group at two time points and, given multiple comparisons, some significant findings may be due to chance. Despite these limitations, the significant main effects suggest that both genotype and intervention group influenced participants’ vegetable consumption frequency (Figure 4). Future studies with larger sample sizes and more participants per phenotype and intervention group at each time point should be powered to identify additional three-way statistical interactions.

The T2R gene family represents a collection of 28 genes found on chromosomes 5, 7, and 12 [26,38] that are expressed in taste bud cells. Given the ability of people to distinguish more distinct bitter tasting compounds than the number of receptors suggests T2R receptors likely respond to more than one bitter ligand [39]. We expanded our SNP-level analysis to cover 20 T2R genes to look for other taste receptors that may provide some insight into the phenotype of our HHL participants. Although our results at the individual SNP level in other T2R genes did not identify associations to changes in vegetable intake within our intervention (Table 5), our multivariate analysis (Figure 5) did identify other loci other than *TAS2R38* that should be considered in future studies, including *TAS2R50* that recognizes the naturally occurring bitter compounds amarogentin and andrographolide [40], and *TAS2R20*, a receptor with no known natural ligand [41]. Within the CAU group our analysis identified SNPs from an additional locus containing three genes in chromosome 7, recently identified as having long-range haptotype structure with *TAS2R38* [42] that contains two receptors with undefined natural ligands, *TAS2R3* and *TAS2R5* [41], and *TAS2R4*, a known receptor for quinine [43].

Given the American Heart Association recommends individual focused interventions for increasing fruit and vegetable intake [44], our findings raise several important issues regarding how we can develop precision medicine approaches in the context of taste perception to inform dietary interventions for heart health. Measuring consumption of specific vegetables that contain glucosinolates and isothiocyanates (e.g., collard greens, broccoli, Brussels sprouts, kale), as well as vegetable preparation methods (e.g., cooked, fresh), could yield more robust associations between bitter taste perception alleles and consumption of bitter vegetables. Conducting a qualitative study among bitter tasters who consume vegetables to learn how and why they have overcome a genetic predisposition to perceive compounds in vegetables as bitter may yield strategies for interventions aiming to increase vegetable consumption. Future research could test whether personalizing diets to specific genetic-based taste profiles increases consumption of specific healthy foods more than generalized dietary advice. Supportive of this concept, a meta-analysis of behavioral interventions found that tailored nutrition interventions aiming to increase fruit and vegetable consumption were more successful than untailored interventions [45,46].

Nutrigenomics and other approaches to tailor nutrition advice and interventions based on genetic and metabolic profiles are increasing as scientists overcome technological and data challenges [47]. In one study, genes associated with energy metabolism were used to personalize a low glycemic index weight management program informed by the Mediterranean diet for participants [48]. The authors observed greater diet adherence to the genetically tailored diets, as well as longer-term reductions in BMI and improved blood glucose levels compared to participants who received a low glycemic index weight management program informed by the Mediterranean diet that was not genetically-tailored [48]. A recent review of nutrigenomic studies did not report any studies that used genes associated with taste perception to inform dietary intervention strategies [47]. Recognizing the important influence that taste perception has on diet and tailoring dietary interventions using this information may be a strategy for engaging participants and improving dietary intervention outcomes.

Reducing heart health disparities requires attention to the many factors driving the disparities. Despite high prevalence of cardiovascular disease among African Americans, this population is under-represented in GWAS studies [49]. Likely explanations include mistrust between African American community members and researchers due to the legacy of unethical medical and genetic studies [50], and imbalances in information and power [51], as well as persistent biases that influence research participation [52]. A strength of the HHL study was our community-based participatory research (CBPR) approach where we worked with a community advisory board, held focus groups with community members, and hired and trained community members as study staff [53,54]. We believe these activities helped build trust between researchers and community-based participants, and helped the research team better understand and meet the expectations that community members had regarding their participation in the genomics portion of this study. Moreover, these activities likely contributed to the high enrollment of African Americans in the genomics arm of the HHL study. In addition to the genomics and lifestyle counseling components of the study, HHL sought to address heart health disparities by increasing access to healthy foods, promoting knowledge of heart healthy choices through a collaboration with local restaurants that included information on healthful menu items and a coordinated monthly newspaper column with information on healthy eating [55], and enhancing clinical care for hypertension in the Lenoir community [23,53]. These strategies were designed to address behavioral and environmental factors that drive heart health disparities in a rural NC population. Combining precision medicine insights to engage participants with CBPR principles and public health strategies that shape the context in which individuals live, work, and play may be a promising approach for reducing cardiovascular health disparities in the US.

This study demonstrates a concept that genes associated with bitter taste perception can influence frequency of vegetable intake in the context of a dietary intervention in a diverse, community-based study sample. The variability in frequency of intake according to participants’ bitter taste perception phenotype could help explain why dietary change interventions report mixed results. Taste has a strong influence over individuals’ dietary habits and should be considered when designing dietary change interventions and in developing novel precision medicine approaches to lifestyle interventions.

## Methods

### The Heart Healthy Lenoir (HHL) Project Overview

The overall goal of the HHL Project was to reduce Cardiovascular Disease (CVD) risk and disparities in CVD risk among Lenoir County, North Carolina residents, as previously described [53,56]. It was conducted in Lenoir County because of its location in the “stroke belt” [57] of eastern North Carolina, where rates of CVD are higher than state and national averages [58] and because it has a large minority population (40% African American) that experiences disproportionally higher rates of CVD [59]. The overall Project included three coordinated studies: a lifestyle intervention study focusing on diet and physical activity [22] a study to improve high blood pressure management at local clinical practices [23] and a study examining associations between genetic markers and change in CVD risk factors. The Project was designed and conducted with input from a local Community Advisory Committee and approved and monitored by the University of North Carolina at Chapel Hill’s Institutional Review Board, with data collected from September 20, 2011 to November 7, 2014 and analyzed in 2017. This trial is registered as # NCT01433484 at clinicaltrials.gov.

### Heart Healthy Lenoir (HHL) Interventions

Participants in the HHL Project (N = 664 in total) could take part in the lifestyle study (N = 339), the high blood pressure study (N = 525) or both (N = 200). All participants were invited to take part in the genomics study. We utilized the data collected at baseline and at the 6-month follow-up that included participants with complete data for the variables of interest in this study, including bitter taste perception phenotype characterized by three SNPs on the *TAS2R38* gene, vegetable intake frequency, and model covariates (N = 497). Twelve participants of the 509 genotyped (2%) were missing data (other than household income) and therefore removed from the analysis. The lifestyle intervention is described in detail elsewhere [22]. Briefly, during the first 6 months, the dietary component of this intervention included four counseling sessions that focused on improving dietary fat and carbohydrate quality, consistent with a Mediterranean dietary pattern. The primary focus of the second counseling session was on increasing fruit and vegetable consumption with a goal of seven total servings per day. The high blood pressure intervention is also described in detail elsewhere [23,53]. Participants in the high blood pressure study received limited dietary counseling by phone, with only 13 receiving a counseling phone call before the 6-month follow-up measurement visit. Accordingly, in this paper, the dietary intervention given to lifestyle study participants is considered the “enhanced” intervention, while the intervention given to those who only participated in the high blood pressure study is considered the “minimal” intervention.

### Genotyping procedure

SNP status was obtained from 505 HHL participants at baseline via DNA isolated from peripheral blood cells using the Infinium Human Omni Express Exome+ BeadChip (Illumina). Genotypes were generated from genomic DNA using the Infinium workflow essentially as described by the manufacturer. DNA was amplified, fragmented, precipitated with isopropanol, and resuspended prior to hybridization onto BeadChips containing 50mer probes. After hybridization, enzymatic single base extension with fluorescently labeled nucleotides was conducted to distinguish alleles. Hybridized BeadChips were imaged using an Illumina iScan to determine intensities for each probe. Corresponding genotypes were extracted from intensity data and called using a standard cluster file within Illumina Genome Studio software. A MAIME-compliant dataset of the microarray data generated is available at the NCBI database of Genotypes and Phenotypes (dbGaP, study ID phs001471).

### Imputing SNPs

All DNA samples identified as either African American (AA, N = 304) or Caucasian American (CAU, N = 201) were imputed for a total of 505 samples. The array data were exported into plink format converted into chromosome-specific variant call format, applying the following filters: merge replicate probes, switch the alternate (ALT) or reference (REF) sequence if deemed necessary by reference, exclude markers where neither REF nor ALT matches the reference, exclude markers where REF is not AGCT. Additionally, in preparation for imputing the following filters were further applied: remove markers not in the reference, fill ALT values in from reference where genotype is entirely homozygous for reference. Samples were imputed twice, once with the Michigan imputation server [60] and once with Beagle (v4.1) [61]. All 505 samples imputed with Beagle were run against the 2504 sample reference panel from 1000 genomes. The Haplotype Reference Consortium (HRC, 65k haplotypes) reference panel was used to run the CAU samples on the Michigan imputation server, and the Consortium on Asthma among African-ancestry Populations in the Americas (CAAPA) reference panel was used to run the AA samples on the imputation server. A brief summary of coverage regarding the panels and how they performed with the target marker set (the markers from the genotyping array) is provided (**Supplemental Table 1**). However, the Illumina genotyping arrays are sparse compared to the reference panels. We filtered our array data for conformity and the markers remaining used for the variant call formatted files (VCF) are indicated (**Supplemental Table 2**).

### *Phased genotype, haplotype, and diplotype* analysis

The phased genotyping data on chromosome 7 for the three *TAS2R38* SNPs (rs713598, rs10246939, and rs1726866) were used to extract the haplotypes of each study subject using the public server at usegalaxy.org [62] to analyze the data with the VCFgenotype-to-hapoltype tool (v1.0.0). VCFtools (v0.1.15) was used to generate all genotype and haplotype frequencies as well as the linkage disequilibrium analyses [63]. The resulting diplotype consisting of the three substitution mutations was used to determine the bitter taste sensitivity phenotype using previously published PROP taste responsiveness with a single PAV haplotype conferring bitter taste [18].

### Outcome variable

We used the Block Fruit and Vegetable Screener [64] to assess vegetable consumption in two mutually exclusive categories: green salads and other types of vegetables. The Block F&V screener is valid for assessing high and low vegetable intake and has been used in African American and White populations [64,65]. Frequency scores were calculated by adding the frequency categories (0 = less than once/week; 1 = once/week; 2 = 2-3 times/week; 3 = 4-6 times/week; 4 = once/day; 5 = 2 or more/day) for the two questions. Frequency scores ranged from 1-10. A score of four is equivalent to about one serving of vegetables per day and a score of five is equivalent to two or more servings per day.

### Covariates

The following covariates were included in the models: sex, age, household income, education, and current smoking status. Taste perception diminishes with age [66] and females are typically more taste sensitive than males [67]. Smoking reduces taste perception [68]. Race, income, smoking status, and education levels are associated with vegetable consumption [69–71]. Sex, smoking status (currently smoking, non-smoker), race (African American or Caucasian), household income (reported in $5,000 incremental categories), and highest year of education achieved, were included as categorical variables. Income was defined as total combined income of participants’ household in the past year, including income from all sources such as wages, salaries, Social Security or retirement benefits, and help from relatives. The mean household income was imputed when data were missing (Table 1). Age was used as a continuous variable.

### Statistical analysis

We used mixed effects models with repeated measures and STATA’s margins command to estimate the adjusted predicted vegetable consumption score for participants within each intervention group and phenotype group at baseline and 6-months follow up. We tested two-way interactions (phenotype group: intervention group and phenotype group: time) and a three-way interaction (phenotype group: intervention: time). Adjusted predicted margins estimate the means for each group of interest, adjusting for the covariates in the mixed effects models [72]. Predicted margins for vegetable consumption scores were contrasted to test whether there were significant differences between participants by intervention group and phenotype group over time. Statistical significance was defined as *p* ≤ 0.05. Statistical analyses were conducted in STATA 15.0 [73]. Principal components analysis and the *p* value of individual SNPs or the SNP: time interaction using mixed effects models with repeated measures was conducted in JMP Pro (v13.2.0, SAS).

## Acknowledgments

This research was supported by National Institutes of Health (NIH) grant 5P50 HL105184 to The University of North Carolina Center for Health Promotion and Disease Prevention (HPDP) with subcontract to the Brody School of Medicine, East Carolina University. This work was conducted at the Center for Health Promotion and Disease Prevention, a Centers for Disease Control and Prevention (CDC) Prevention Research Center (5U48DP001944), the Center for Health Equity Research, and the McAllister Heart Institute at The University of North at Chapel Hill. We are grateful to multiple administrative teams that supported this work, and to Dr. Steven Ziesel for his input in the early stages of this work. We give special thanks to our Lenoir County Community Advisory Committee who provided helpful guidance with this project and to our study participants, whose willing participation made this study possible.

